# FISH-quant v2: a scalable and modular analysis tool for smFISH image analysis

**DOI:** 10.1101/2021.07.20.453024

**Authors:** Arthur Imbert, Wei Ouyang, Adham Safieddine, Emeline Coleno, Christophe Zimmer, Edouard Bertrand, Thomas Walter, Florian Mueller

## Abstract

Regulation of RNA abundance and localization is a key step in gene expression control. Single-molecule RNA fluorescence in-situ hybridization (smFISH) is a widely used single-cell-single-molecule imaging technique enabling a quantitative understanding of gene expression and its regulatory mechanisms. Recent progress in experimental techniques provides larger data-sets, requiring adequate tools for data analysis and exploration. Here, we present FISH-quant v2, a highly modular analysis tool accessible both for non-experts and experts, which we validated and applied on large-scale smFISH image datasets. Our package allows the user to detect isolated and clustered mRNA spots, segment nuclei and cells, quantify RNA localization patterns and visualize these results at the single-cell level.

## Introduction

Regulation of gene expression is essential for a cell to fulfill its basic functions, and its dysregulation can lead to serious failures at the cellular, tissular and organism levels. Transcription levels are not only tightly regulated, but for many genes it has now been demonstrated that their transcripts accumulate in specific regions in the cell, thereby producing intricate localization patterns. Such subcellular targeting of mRNAs is thought to play an important role for the spatial control of gene expression; in which improper RNA trafficking is linked to an increasing number of diseases (1, 2). However, the function and mechanisms of RNA localization are not fully understood and we lack a view of this process at the transcriptomic scale.

RNA abundance and localization can be studied at a large scale by image-based assays, where individual mRNA molecules are visualized by single molecule Fluorescence in situ hybridization (smFISH). smFISH allows for the visualization of individual mRNA molecules in their native cellular environment (3, 4) by targeting each mRNA with several fluorescently labeled oligonucleotides. Many variants of this method exist, with optimizations regarding signal to noise ratio (SNR), experimental protocol, targeting specificity, scalability, automatization and cost. Furthermore, an increasing number of multiplexing methods have also been proposed over the last years, enabling the simultaneous imaging of a large number of different RNAs (5, 6) in cells and tissues. Usually, smFISH experiments are complemented by the use of one or several fluorescent markers highlighting relevant compartments in the cell, such as the nucleus, the cytoplasm or any organelle that might serve as a reference, depending on the focus of the study.

These scalable imaging techniques produce extremely large and complex image data sets exploring spatial aspects of large portions of the transcriptome. While large-scale imaging methods provide a systematic tool to understand RNA localization at a systems level, they come at a price: the need of fully automated, robust image analysis and user-friendly software tools to analyze such data sets and to fully exploit their potential (7, 8).

Several specifications can be defined *a priori* for such an analysis tool. It should be simple enough to be mastered by non-experts, especially non-coders. Yet, it should be flexible enough to address different experimental designs and aims and be adapted from a common algorithmic backbone. With the same modules, users should be able to both perform a high content screening analysis in a remote cluster, and a local analysis over a single image. Finally, the software should integrate the latest generation of computer vision algorithms, especially those from the machine learning community that have been shown to provide excellent results, in particular deep learning based methods for image segmentation (9–11).

Here, we introduce a Python-based version of our widely adopted software package FISH-quant (12) for the analysis of smFISH images. Contrary to FISH-quant v1 in Matlab version, we address and improve on each of the specifications mentioned above. The switch to Python allows us to develop a flexible, free and fully open-source software. FISH-quant v2 enjoys a better integration to other open source tools and frameworks, from data analysis to web-based user interaction. Last but not least, FISH-quant v2 facilitates the use of machine learning or deep learning algorithms with the import of dedicated packages, such as scikit-learn(13) or tensorflow^13^. We also improve the scalability and the modularity of the package: the software has now been applied to several High Content Screening projects ^14–16^. Lastly, by using ImJoy (18), a recently developed data analysis framework, we provide web-based graphical user interfaces (GUI) for both launching image analysis and downstream analysis of the results and the computation can be performed locally or seamlessly scale to powerful remote computing servers.

## Materials and Methods

### Big-FISH: the core analysis package

The repository Big-FISH contains the entire Python code used for the actual analysis. It is organized in several subpackages performing dedicated steps:

- I/O operations, images preprocessing and results postprocessing (*bigfish.stack*)
- mRNA spot detection (*bigfish.detection*)
- nucleus and cell segmentation (*bigfish.segmentation*)
- feature computation and coordinates analysis (*bigfish.classification*)
- Plotting of results (*bigfish.plot*) and
- Application of deep learning algorithms for segmentation (*bigfish.deep_learning*).

Dependencies are limited to standard Python scientific libraries: scientific computing (numpy (19), scipy (20)), data wrangling (pandas (21)), image analysis (scikit-image (22)), visualization (matplotlib (23)) and machine learning (scikit-learn (13), tensorflow (24)).

The GitHub repository is using continuous integration providing increased robustness of the released code, through unitary testing, version control and automatically up-to-date documentation. The package is hosted under a BSD 3-Clause License.

### Example datasets

Two datasets were used for the development and validation of FISH-quant. First, from a screen studying local translation, consisting of 526 fields of view (Dapi and smFISH channels) from 57 separate experiments (27 different mRNAs under different experimental conditions)(16). 3D images with a z-spacing of 0.3 μm were acquired on two different systems: (i) a Zeiss AxioimagerZ1 wide-field microscope equipped with a motorized stage, a camera scMOS ZYLA 4.2 MP, using 63x and 100x oil objectives, (ii) Nikon Ti fluorescence microscope equipped with ORCA-Flash 4.0 digital camera (HAMAMATSU). Second, from a screen focusing on local translation of centrosomal mRNAs. The data-set consisted of 3678 fields of view (Dapi, smFISH, CellMask and GFP channels) from 218 experiments (15). 3D images were acquired with an automated spinning disk microscope (Opera, Perkin Elmer), equipped with a 63x water objective. Z-spacing was 0.3 μm.

## Results

The analysis of smFISH images aims at counting and localizing individual RNAs with respect to single cells and other cellular landmarks. It typically encompasses a number of steps: (1) segmenting cells and the relevant cellular compartments such as nuclei (depending on the focus of the study and the markers employed), (2) detecting individual RNA molecules appearing as diffraction-limited spots, (3) assignment of spots to cells and (4) analysis of expression levels and RNA localization patterns (4, 12, 25–27), potentially in combination with other phenotypic features (15, 28).

We designed FISH-quant v2 to match this workflow in a flexible and efficient way. This version is entirely open-source and hosted on GitHub under the FISH-quant organization (**Figure 1,** https://github.com/FISH-quant). It is organized in several resources with dedicated repositories and documentation. First, a Python package (Big-FISH) providing the core code for performing computation and analysis. Second, detailed interactive examples with test data for each analysis step implemented in Jupyter notebooks. These examples can be run directly on Binder (29), a free and reproducible Jupyter notebook service, without local installation. Third, ImJoy plugins(18) provide a graphical user-interface for the most commonly used workflows, and an interactive tutorial that can also run directly without local installation. Lastly, code from future projects either using or further improving FISH-quant will be also hosted here, creating a valuable, centralized resource for the community. A landing page (https://fish-quant.github.io/) directs new users to the most relevant resource for their analysis needs.

**Figure 1.**
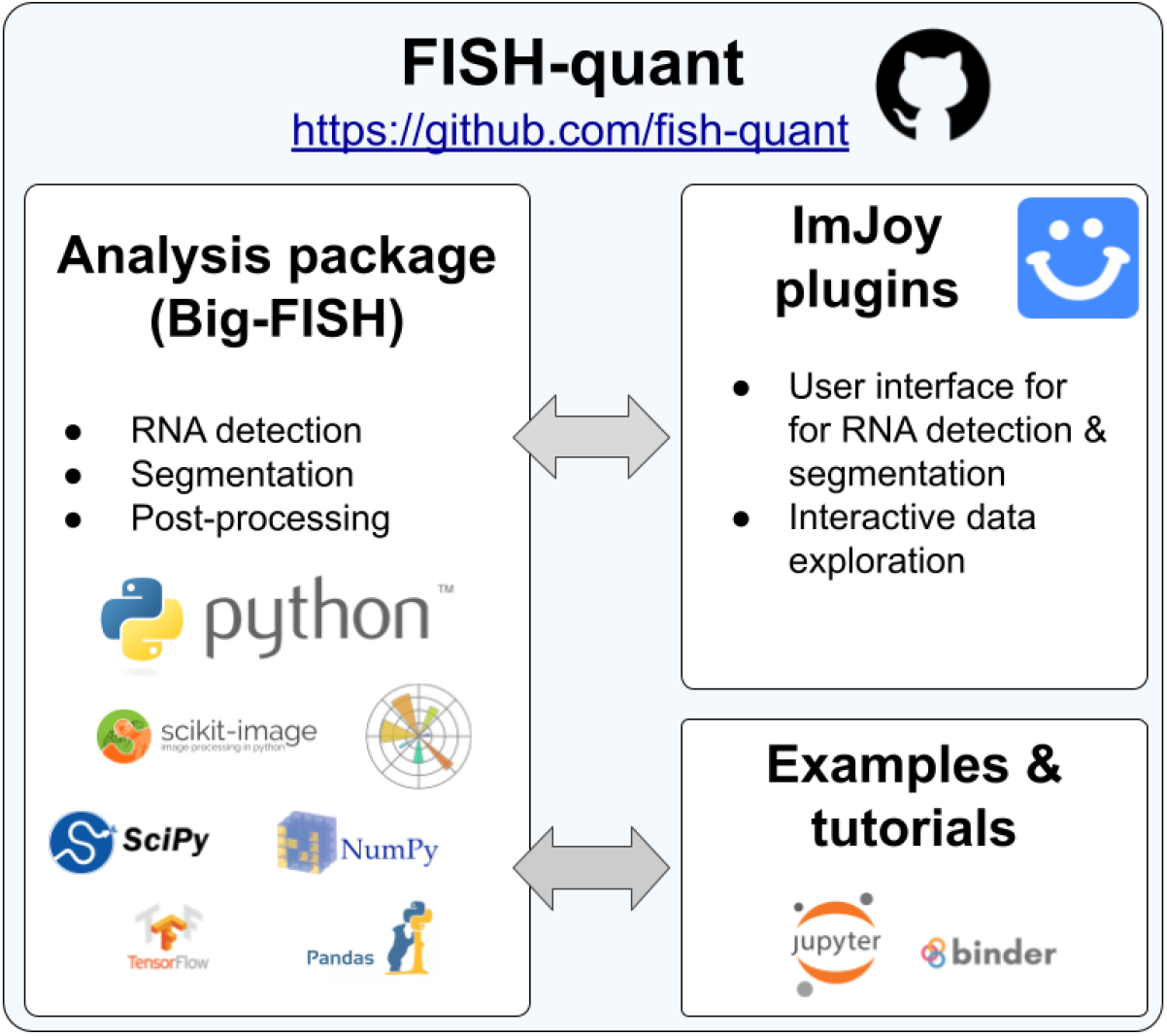
Organization of FISH-quant. FISH-quant is hosted on Github and consists of several interconnected repositories. The Python core package contains the entire analysis code, which is used by both the ImJoy plugins and the example and tutorial repository.

### Python core analysis package

We choose Python for the implementation of the core analysis package for several reasons: it allows the development of a free and fully open-source software, it provides established libraries for data and image analysis and is the language of choice for deep-learning implementations. Lastly, it can be interfaced with other tools and frameworks, from data analysis to web design, for instance with ImJoy(18) to provide interactive tools for user interaction and data inspection.

Our Python package includes several independent subpackages fitting the described workflow (see Material and Methods for more details): preprocessing, segmentation, detection, and analysis. We designed each subpackages with clearly defined input and output data formats, which will be automatically checked. This then allows using each of these packages independently in a modular fashion. Users can thus create a customized analysis workflow, starting with pre-processing of images to statistical results. This modular design also permits to easily integrate external methods, for instance, a new segmentation method can be combined with our spot detection algorithm.

Here, we will only provide an overview of these submodules (**Figure 2**). For a more detailed description of algorithms and methods, we refer to the detailed documentation (https://big-fish.readthedocs.io/en/stable/) and the dedicated tutorials (https://github.com/fish-quant/big-fish-examples). These tutorials can be run directly in the browser with provided test data, and thus allow new users to immediately test these tools. The described methods were developed and validated with the data from two large-screen smFISH studies (15, 16) (see Material and Methods).

**Figure 2.**
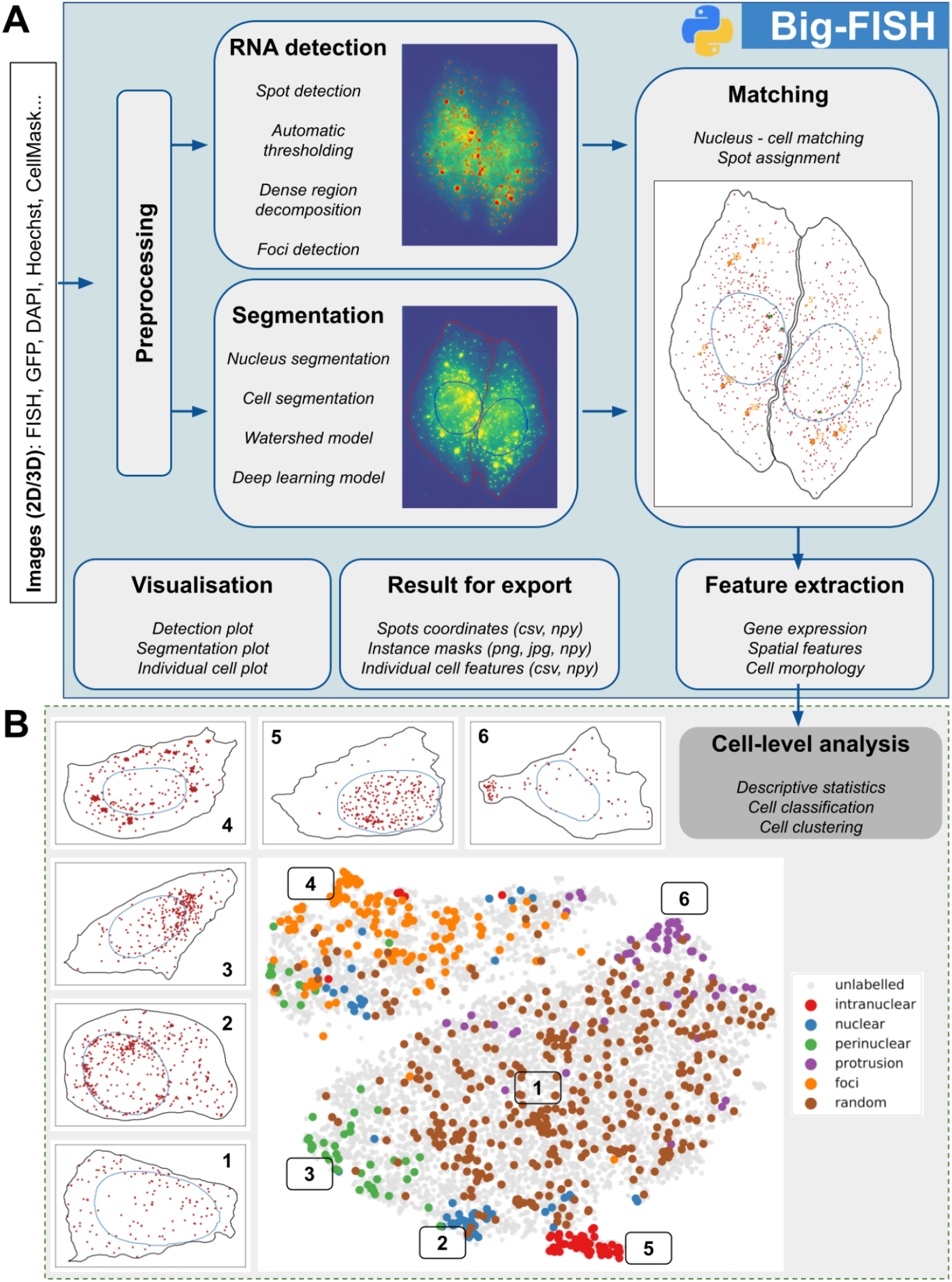
Big-FISH: the core analysis Python analysis package. **(A)** Main modules illustrated with a typical analysis workflow. Shown are also the inputs and outputs that are created at the different steps. (**B**) As a final result of the analysis of big-FISH, each cell is described with a set of features reflecting RNA abundance and localization. These features can then be used to perform analysis on the cell population. Shown are results from our RNA localization screen where cells are grouped based on their RNA localization pattern(16). The t-SNE plot projects 15 localization features for smFISH experiments against 27 different genes. Each dot is one cell. The color-coded dots are manual annotations of 6 different localization patterns. Images are examples of individual cells displaying a typical localization pattern of this region of the t-SNE plot.

The visualization module provides the functionality to visualize the results of each intermediate step in the analysis workflow, and thus provides valuable visual quality control.

For image handling and preprocessing, we implemented a number of different utility functions to read, write, normalize, cast, filter, and project images. Different image file formats are natively supported and 2D and 3D images can be processed.

The detection module implements the methods required to detect spots in 2D or 3D images (**Figure 2**). While initially designed to detect individual mRNAs, the same methods can also be used to detect other spot-like structures (15), such as centrosomes, P-bodies, etc. Strong local accumulation of RNAs, e.g. at sites of local translation(16), can lead to an underdetection. For such cases, we provide tools to decompose these dense regions and estimate the number of spots based on our earlier work (see **Supplementary Note 1**) (27). An important aspect of the detection subpackage is the ability to detect spots without setting any pixel intensity threshold (see **Supplementary Note 1)**. This parameter can be automatically inferred from the image. Such automatization eliminates human intervention and allows scaling to large datasets, such that the subpackage can process thousands of images.

The segmentation module contains several algorithms and utility functions for segmentation and postprocessing. It provides deep-learning-based approaches (**Figure 2, Supplementary Note 2**), and postprocessing steps to improve segmentation accuracy. Each segmented instance can then be refined and cleaned, and morphological properties can be computed for these components (**Supplementary Note 3**).

The cell matching module allows combining results from detection and segmentation permitting to analyze RNA abundance and distribution at the single-cell level. Every detected spot can be assigned to a specific area of interest, for instance, a cell or a nucleus. mRNA expression levels are extracted within this module, as this is usually the minimum information we would like to extract from this kind of images.

The localization feature extraction module permits the extraction of further information to study the spatial distribution of mRNA molecules. It gathers methods to format spot positions and coordinates of cellular landmarks and compute several spatial features at the single-cell level (**Figure 2, Supplementary Note 3**). These features allow a statistical description of the cell population (15, 17) or can feed a classification model permitting to classify individual cells based on their RNA localization patterns (16) (**Figure 2**).

### Interactive User Interface with ImJoy

Our Python core analysis package provides flexibility and scalability since these modules can be adapted to the specific analysis need of a given project. However, they require at least a minimum knowledge of Python to establish a complete workflow by using the provided tutorials.

To provide simple access for users with no computational background, we implemented several plugins with graphical user interfaces for our computational platform ImJoy (18). These plugins provide the most commonly used analysis workflow, as we determined from the usage of the Matlab version of FISH-quant, and will thus be suited for a large number of user cases (**Figure 3**). First, a plugin to perform deep-learning-based segmentation. This is currently built on top of CellPose (11), but thanks to our modular design, this can be easily exchanged if more performant methods are available in the future. Second, detection of both isolated and clustered RNA. Detection results can be conveniently inspected with Kaibu image viewer plugin in ImJoy and different detection settings interactively investigated. Batch processing of entire folders is also possible. Lastly, detection results can be assigned to segmented cells and nuclei. We provide an interactive demo version of this plugin that can run directly in the browser without any local installation (https://fish-quant.github.io/fq-interactive-docs/#/fq-imjoy).

**Figure 3.**
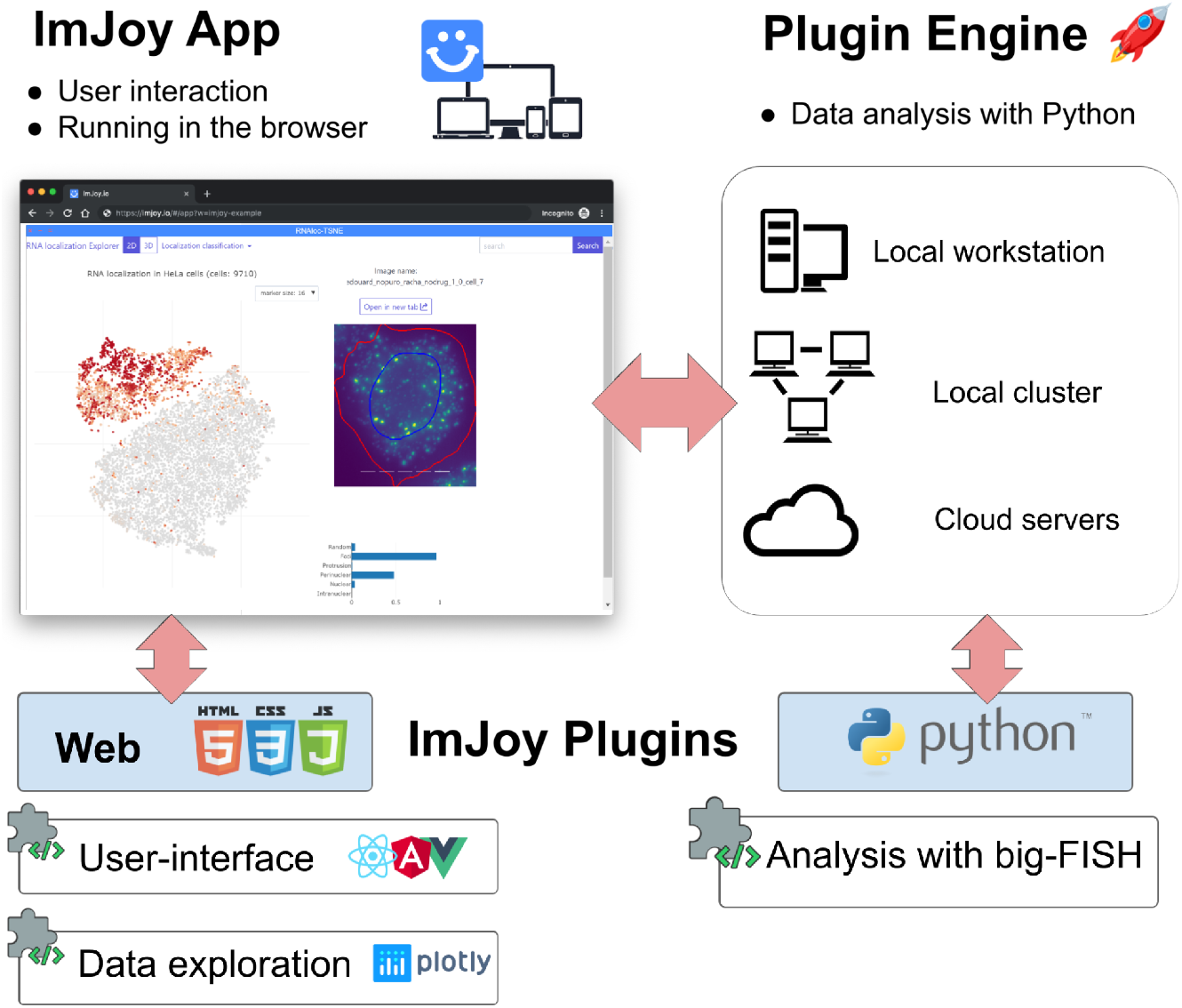
ImJoy. Schematic view of Imjoy’s architecture. ImJoy’s core is a Progressive Web App whose functionalities are provided by plugins that can be written in different programming languages. ImJoy can perform computations in the browser (including offline), locally or remotely via plugin engines.

Using ImJoy provides several advantages beyond simply providing a user interface. Due to its distributed design that separates GUI from computation plugins, it natively allows user-friendly remote computing which allows access to massive data storage and powerful computation resources including GPUs. ImJoy’s is a browser-based app where the user-interface plugin is implemented with JavaScript/CSS/HTML. ImJoy then transparently calls the computation functions in the BigFISH package running on a Python plugin engine (e.g. Jupyter server) to perform the actual smFISH analysis task (**Figure 3**). While this plugin can run on the local workstation, it can be located in a computational cluster even in the cloud or seamlessly switching between them. As illustrated by the demo version, where the engine is running on Binder(29). Once the plugin engine is installed on the remote resource, the end-user can connect with ImJoy and will be confronted with the same interface, independently on where the analysis is actually performed. Interestingly, this front-end interface can also be opened with mobile devices, providing easy access.

ImJoy plugins implemented in JavaScript can not only provide modern and reactive user-interfaces, but also profit from the extensive JavaScript data visualization libraries to build interactive data-inspection tools. Such interactivity is becoming increasingly important, especially when large and complex data-sets are analyzed where static plots are too limited. As a case example, we provide an interactive t-SNE plot for the data shown in Figure 2 (https://fish-quant.github.io/fq-interactive-docs/#/rnaloc-tsne). This plugin can be run without local installation, and enables the user to explore and interact with these complex data.

## Discussion

Here, we present FISH-quant v2, a Python-based software for the analysis of smFISH images. It is built around a core-analysis package, implemented following rigorous software development guidelines, with detailed interactive documentation and tutorials. This package consists of several interchangeable modules permitting the construction of highly flexible workflows for specific analysis needs. For standard workflows, we also provide user interfaces in ImJoy accessible to biologists without programming skills, which can be used locally or scaled to larger remote computational resources. As demonstrated in two recently published studies (15, 16), FISH-quant can be used for large screening data sets thanks to its scalability. Spot detection, segmentation, feature engineering and pattern recognition can be performed over thousands of cells without fine-tuning parameters for every image.

We designed FISH-quant v2 based on the successful previous implementation in Matlab (12) integrating new features and user feedback we obtained from several projects over several years. The entire core package is written in Python since this allowed us to address the above-mentioned requirements for a smFISH analysis tool. We use established scientific libraries (see Methods), and kept these dependencies to a minimum facilitating installation, maintenance and the integration with other analysis frameworks. These libraries are developed, validated and maintained by a large scientific community, ensuring long-term support and availability. We further use strict version control, guaranteeing reproducibility. Lastly, all dependencies, as well as FISH-quant, are open-source, thus can be used free of charge, both on local and remote computational infrastructures, and thus analysis can easily be scaled to larger data volumes.

The organization of the analysis modules in the core package matches key steps in smFISH image analysis, but with a special focus on flexibility. All modules (Preprocessing, RNA detection, segmentation as well as data inspection and analysis) can be run independently or replaced by external code, by respecting a strict data format. This allows FISH-quant to be adapted to the respective analysis needs, and build custom workflows.

While this flexibility is important, many users require a standard workflow and don’t have programming experience. For these cases, we provide ImJoy plugins with a convenient user interface running in the browser(18). These interfaces are built with modern web libraries and are thus intuitive, and no experience in Python is required to analyze data. Lastly, these ImJoy plugins can be readily extended by more experienced users to further adapt them to their needs. A detailed documentation and interactive tutorial further help new users to get started quickly.

In summary, we present with FISH-quant v2 a rigorously validated analysis platform for smFISH data, developed to match the analysis requirements of large data-sets. Its modularity permits analysis tasks ranging from small data-sets with the help of a graphical user-interface to large-scale screens requiring computational clusters.

## Availability

The entire code for the analysis described in this paper is available on GitHub: https://github.com/fish-quant

This study includes no data deposited in external repositories.

## References

1. Buxbaum,A.R., Haimovich,G. and Singer,R.H. (2014) In the right place at the right time: visualizing and understanding mRNA localization. Nat. Rev. Mol. Cell Biol., 16, 95–109.

2. Chin,A. and Lécuyer,E. (2017) RNA localization: Making its way to the center stage. Biochim. Biophys. Acta BBA - Gen. Subj., 1861, 2956–2970.

3. Raj,A., van den Bogaard,P., Rifkin,S.A., van Oudenaarden,A. and Tyagi,S. (2008) Imaging individual mRNA molecules using multiple singly labeled probes. Nat. Methods, 5, 877–879.

4. Tsanov,N., Samacoits,A., Chouaib,R., Traboulsi,A.M., Gostan,T., Weber,C., Zimmer,C., Zibara,K., Walter,T., Peter,M., et al. (2016) SmiFISH and FISH-quant - A flexible single RNA detection approach with super-resolution capability. Nucleic Acids Res., 44.

5. Moffitt,J.R. and Zhuang,X. (2016) RNA Imaging with Multiplexed Error-Robust Fluorescence In Situ Hybridization (MERFISH). Methods Enzymol., 572, 1–49.

6. Eng,C.-H.L., Lawson,M., Zhu,Q., Dries,R., Koulena,N., Takei,Y., Yun,J., Cronin,C., Karp,C., Yuan,G.-C., et al. (2019) Transcriptome-scale super-resolved imaging in tissues by RNA seqFISH. Nature, 568, 235–239.

7. Das,S., Vera,M., Gandin,V., Singer,R.H. and Tutucci,E. (2021) Intracellular mRNA transport and localized translation. Nat. Rev. Mol. Cell Biol., 10.1038/s41580-021-00356-8.

8. Pichon,X., Lagha,M., Mueller,F. and Bertrand,E. (2018) A Growing Toolbox to Image Gene Expression in Single Cells: Sensitive Approaches for Demanding Challenges. Mol. Cell, 71, 468–480.

9. Ronneberger,O., Fischer,P. and Brox,T. (2015) U-Net: Convolutional Networks for Biomedical Image Segmentation. In Navab,N., Hornegger,J., Wells,W.M., Frangi,A.F. (eds), Medical Image Computing and Computer-Assisted Intervention – MICCAI 2015. Springer International Publishing, pp. 234–241.

10. Falk,T., Mai,D., Bensch,R., Çiçek,Ö., Abdulkadir,A., Marrakchi,Y., Böhm,A., Deubner,J., Jäckel,Z., Seiwald,K., et al. (2019) U-Net: deep learning for cell counting, detection, and morphometry. Nat. Methods, 16, 67–70.

11. Stringer,C., Wang,T., Michaelos,M. and Pachitariu,M. (2021) Cellpose: a generalist algorithm for cellular segmentation. Nat. Methods, 18, 100–106.

12. Mueller,F., Senecal,A., Tantale,K., Marie-Nelly,H., Ly,N., Collin,O., Basyuk,E., Bertrand,E., Darzacq,X. and Zimmer,C. (2013) FISH-quant: automatic counting of transcripts in 3D FISH images. Nat. Methods, 10, 277–278.

13. Pedregosa,F., Varoquaux,G., Gramfort,A., Michel,V., Thirion,B., Grisel,O., Blondel,M., Prettenhofer,P., Weiss,R., Dubourg,V., et al. (2011) Scikit-learn: Machine Learning in Python. J. Mach. Learn. Res., 12, 2825–2830.

14. Abadi,M., Agarwal,A., Barham,P., Brevdo,E., Chen,Z., Citro,C., Corrado,G.S., Davis,A., Dean,J., Devin,M., et al. (2015) TensorFlow: Large-scale machine learning on heterogeneous systems.

15. Safieddine,A., Coleno,E., Salloum,S., Imbert,A., Traboulsi,A.-M., Kwon,O.S., Lionneton,F., Georget,V., Robert,M.-C., Gostan,T., et al. (2021) A choreography of centrosomal mRNAs reveals a conserved localization mechanism involving active polysome transport. Nat. Commun., 12, 1352.

16. Chouaib,R., Safieddine,A., Pichon,X., Imbert,A., Kwon,O.S., Samacoits,A., Traboulsi,A.-M., Robert,M.-C., Tsanov,N., Coleno,E., et al. (2020) A Dual Protein-mRNA Localization Screen Reveals Compartmentalized Translation and Widespread Co-translational RNA Targeting. Dev. Cell, 54, 773–791.e5.

17. Pichon,X., Moissoglu,K., Coleno,E., Wang,T., Imbert,A., Peter,M., Chouaib,R., Walter,T., Mueller,F., Zibara,K., et al. (2020) The kinesin KIF1C transports APC-dependent mRNAs to cell protrusions. bioRxiv, 10.1101/2020.11.30.403394.

18. Ouyang,W., Mueller,F., Hjelmare,M., Lundberg,E. and Zimmer,C. (2019) ImJoy: an open-source computational platform for the deep learning era. Nat. Methods, 16, 1199–1200.

19. Harris,C.R., Millman,K.J., van der Walt,S.J., Gommers,R., Virtanen,P., Cournapeau,D., Wieser,E., Taylor,J., Berg,S., Smith,N.J., et al. (2020) Array programming with NumPy. Nature, 585, 357–362.

20. SciPy: Open Source Scientific Tools for Python – ScienceOpen.

21. McKinney,W. (2010) Data Structures for Statistical Computing in Python. Proc. 9th Python Sci. Conf., 10.25080/Majora-92bf1922-00a.

22. van der Walt,S., Schönberger,J.L., Nunez-Iglesias,J., Boulogne,F., Warner,J.D., Yager,N., Gouillart,E. and Yu,T. (2014) scikit-image: image processing in Python. PeerJ, 2, e453.

23. Hunter,J.D. (2007) Matplotlib: A 2D Graphics Environment. Comput. Sci. Eng., 9, 90–95.

24. Abadi,M., Agarwal,A., Barham,P., Brevdo,E., Chen,Z., Citro,C., Corrado,G.S., Davis,A., Dean,J., Devin,M., et al. (2016) TensorFlow: Large-Scale Machine Learning on Heterogeneous Distributed Systems. ArXiv160304467 Cs.

25. Stoeger,T., Battich,N., Herrmann,M.D., Yakimovich,Y. and Pelkmans,L. (2015) Computer vision for image-based transcriptomics. Methods, 85, 44–53.

26. Battich,N., Stoeger,T. and Pelkmans,L. (2013) Image-based transcriptomics in thousands of single human cells at single-molecule resolution. Nat. Methods, 10.

27. Samacoits,A., Chouaib,R., Safieddine,A., Traboulsi,A.-M., Ouyang,W., Zimmer,C., Peter,M., Bertrand,E., Walter,T. and Mueller,F. (2018) A computational framework to study sub-cellular RNA localization. Nat. Commun., 9, 4584.

28. Battich,N., Stoeger,T. and Pelkmans,L. (2015) Control of Transcript Variability in Single Mammalian Cells. Cell, 163, 1596–1610.

29. Jupyter,P., Bussonnier,M., Forde,J., Freeman,J., Granger,B., Head,T., Holdgraf,C., Kelley,K., Nalvarte,G., Osheroff,A., et al. (2018) Binder 2.0 - Reproducible, interactive, sharable environments for science at scale. Proc. 17th Python Sci. Conf., 10.25080/Majora-4af1f417-011.

